# Noninvasive Control of Seizure Threshold with Acoustically Targeted Chemogenetics

**DOI:** 10.1101/2025.05.28.656723

**Authors:** Honghao Li, Shirin Nouraein, Sangsin Lee, Schuyler S. Link, Emma K. Raisely, Jerzy O. Szablowski

## Abstract

Many neurological and psychiatric diseases are characterized by pathological neuronal activity. Current treatments involve drugs, surgeries, and implantable devices to modulate or remove the affected region. However, none of these methods can be simultaneously non-invasive and possess site- and cell type specificity. Here, we apply a non-invasive neuromodulation approach called Acoustically-Targeted Chemogenetics, or **ATAC**, to increase the seizure threshold. Here, the ATAC approach used a multi-point focused ultrasound to transiently open the blood-brain barrier of the whole hippocampus and transduce pyramidal neurons with engineered G-protein-coupled receptors to inhibit their activity. To express the engineered receptors in the mouse hippocampus, we used a recently engineered viral vector optimized for ultrasound-based gene delivery to the brain, AAV.FUS.3. In a mouse flurothyl seizure model, we showed successful gene delivery throughout the hippocampus, a significant neuronal activity inhibition as evidence by an increase in seizure threshold. Finally, we benchmarked these effects against a clinically prescribed drug that acts without spatial precision.

## INTRODUCTION

Neurological and psychiatric diseases are common, affecting one in five U.S. adults, causing a significant personal burden [1-3]. Many prevalent conditions, such as epilepsy, Alzheimer’s disease, and schizophrenia, involve uncontrolled neuronal activation with specific spatial locations, leading to local or global neuronal hyperactivity [4-6]. However, conventional pharmacological treatments for these diseases act throughout the central nervous system (**CNS**), without spatial specificity, activating both the regions of the brain affected by the disease and those that are not, thus causing significant side effects. Surgical interventions, including implantation of devices, can provide spatial specificity but are invasive, causing tissue damage [7, 8]. Moreover, only some of the patients qualify for such treatment based on their individual risks and personal preferences [9]. Therefore, there is an unmet need for one-time non-invasive treatments that can provide spatially precise neuronal control without the need for invasive procedures. Gene therapy is one of the most promising emerging approaches to treating human disease. For example, optogenetics is promising with spatial, cell-type, and temporal control of the treatment, but requires invasive implantation of optical fibers to target deep or large brain regions [10, 11].

Acoustically targeted chemogenetics (**ATAC**) has recently emerged as a promising non-invasive neuromodulation strategy [12] that retains spatial, cell-type, and temporal precision. ATAC integrates focused ultrasound blood-brain barrier opening (**FUS-BBBO**) for precise spatial targeting, adeno-associated virus (**AAV**) vectors for gene delivery to specific cell types, and chemogenetic designer receptors exclusively activated by designer drugs (**DREADDs**) for modulating targeted neurons using clozapine-N-oxide (**CNO**) (**Fig. 1a**). FUS is a well-established biomedical technology that enables precise, non-invasive targeting of deep brain tissues with millimeter spatial accuracy [13, 14]. When combined with microbubbles, FUS can induce localized, temporary, and reversible blood-brain barrier opening [15, 16]. This technology has been successfully applied across various animal species [16, 17] and humans[18-20], enabling the delivery of viral vectors, particularly clinically relevant AAV9, as well as proteins and small molecules to the FUS-insonated site [21-25]. DREADDs offer an approach to reversibly and selectively modulate neuronal activity [26, 27] with hM4Di(**Gi**), which reduces neuronal firing upon binding to a cognate ligand by activating the G protein-gated inwardly rectifying potassium (**GIRK**) channels [26] (**Fig. 1b**). While Clozapine-N-oxide CNO was initially selected as the primary DREADD actuator due to its activation kinetics, recent studies have shown that CNO itself does not effectively cross the blood-brain barrier; instead, subclinical levels of its metabolite, clozapine, is responsible for binding to and activating DREADDs, turning CNO into a prodrug for low-dose clozapine [28, 29]. Multiple studies have shown the effectiveness of suppressing neuronal activity in various animal models of disease by selectively expressing the DREADDs under cell-type specific promoters, especially in excitatory neurons [30-33]. Therefore, DREADDs are being considered for clinical translation [34]. If such therapies could be delivered noninvasively, their applicability would be expanded to a broader range of patients who cannot or choose not to undergo brain surgery. In order to improve the efficiency and specificity FUS-BBBO based gene delivery, we recently developed a novel viral vector based on AAV9, called AAV.FUS.3 [35]. This engineered vector showed 12-15-fold improvement in brain tissue specificity over the liver in two different mouse strains, improved neuronal tropism, and overall transduction efficiency at the brain focus at up to 100-fold lower weight-normalized doses compared to clinically approved systemic AAV9 gene therapy (Zolgensma). Reducing the dose of AAV is desirable as it can reduce the substantial cost of therapy production and potentially reduce systemic side effects [36]. While AAV.FUS.3 showed superior gene delivery over AAV9 after FUS-BBBO, it has not yet been established whether the delivery is efficient enough to exert noninvasive neuromodulation with ATAC, and whether ATAC can modulate a clinically-relevant parameter, such as the seizure threshold, with a large HPC region coverage.

**Figure 1.**
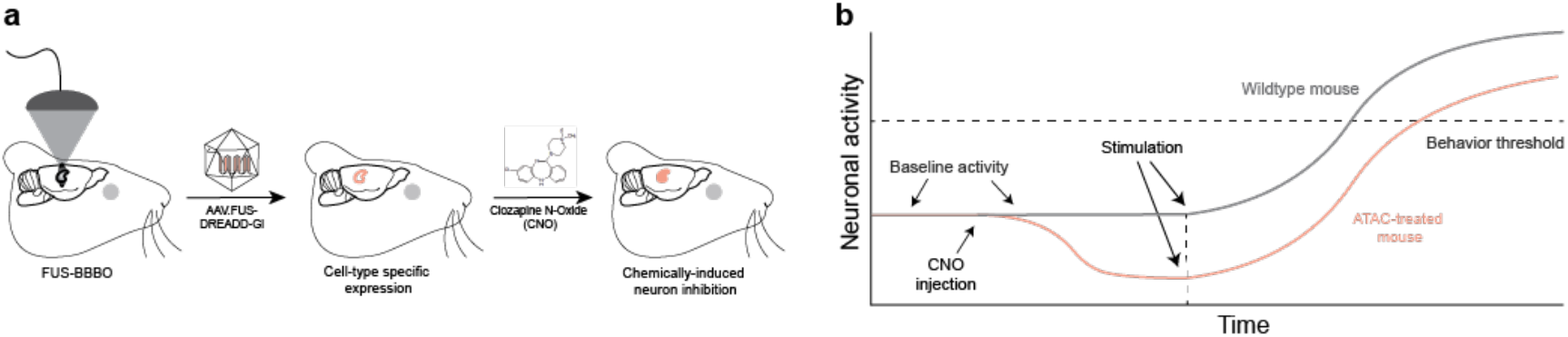
ATAC-AAV.FUS.3 paradigm and neuronal activity comparison. a, To generate an ATAC-treated mouse, the blood-brain barrier (BBB) is locally opened using focused ultrasound (FUS), allowing systemically injected AAV.FUS.3 encoding DREADD-Gi to enter the insonated area. After a few weeks, DREADD-Gi is selectively expressed in the targeted regions with cell-type specificity. The DREADD-Gi-expressing neuron activity can then be inhibited at any desired time using a chemogenetic drug such as CNO. b, Following CNO injections in ATAC mice, basal neuronal activity decreases due to the opening of potassium channels. During stimulation, the ATAC-treated mouse is expected to exhibit a delayed onset of behavioral responses.

Here, we present the state of the art of ATAC, where we cover the entirety of the hippocampus (**HPC**) to increase the seizure threshold in a simple behavioral model. To that end, we used inhalable flurothyl exposure as a means of inducing an acute clonic seizure. This approach provides a robust tool to assess the efficacy of neuromodulation after DREADD delivery with AAV.FUS.3, a recently engineered vector that has not been yet tested in a behavioral model [35]. Furthermore, we benchmark our new ATAC protocol against the clinically approved anti-epileptic drug (**AED**) valproic acid (**VPA**) to evaluate its neuronal suppressive effects, as determined by the delay in the flurothyl-induced seizure threshold. Our findings reveal that ATAC treated mice exhibited a significant delay in seizure onset.

## RESULTS

We first examined the number of FUS-BBBO sites that could be targeted throughout the HPC without causing tissue damage or hemorrhage. Based on our previous study, 4– 6 FUS targeting sites per HPC per hemisphere did not overtly affect survival or behavior of the animal; however, gene delivery efficiency remained inconsistent, and large volume dorsal HPC transduction efficiency was not studied [37]. We targeted the HPC regions that are close to the cortex for dorsal HPC and the center of ventral HPC to avoid the unwanted side transduction of the thalamus, preventing potential vital function inhibition (**Fig. 2a**). To maximize BBB opening efficiency and viral delivery, we designed a protocol targeting 12 sites per hemisphere across two distinct layers in dorso-ventral direction separating each site in medio-lateral and anterior-posteriro directions by the full-width half maximum (**FWHM**) pressure of the beam (**Fig. 2b**). Thus, the center of each targeted site was spaced 0.8mm-1.0mm apart, with the FUS beam insonated vertically from the top of the brain. The order of FUS insonation started from the anterior of the dorsal HPC (**dHPC**) to the ventral HPC (**vHPC**) and back to the dHPC to minimize the transducer travel time between cycles (**Supp Fig. 1a and 1b**). FUS was coupled with an intravenous (**i.v**.) injection of microbubbles (**MB**). We chose the opening parameters based on previous studies [12, 37, 38] and preliminary data in our lab.

**Figure 2.**
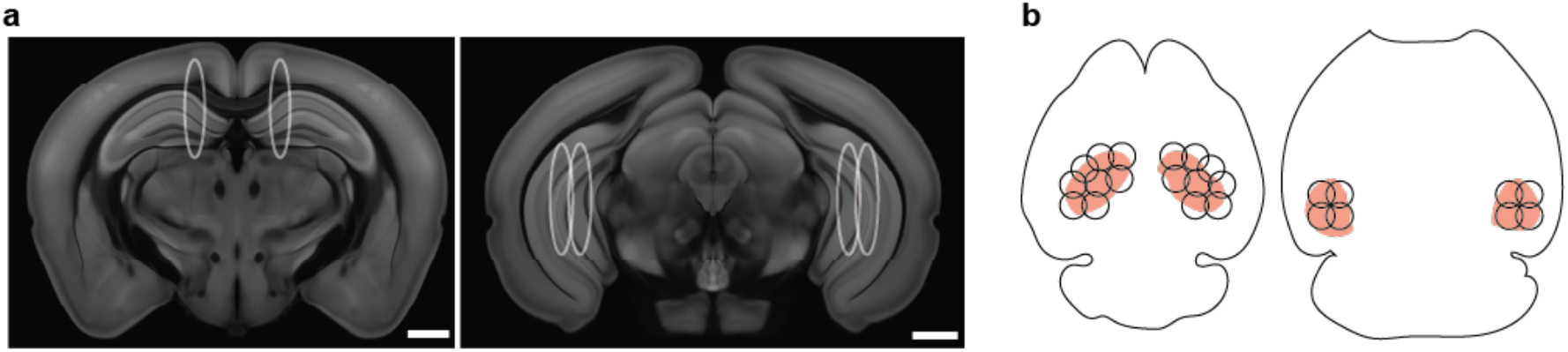
Focused ultrasound (FUS) targeting strategies. **a**, T1-weighted MRI images of the mouse brain coronal atlas obtained from the Allen Institute and an online source. The left image is the dorsal hippocampus, and the right is the ventral hippocampus. The white circles represent the FWHM of pressure of the focused ultrasound beam generated by the transducer according to the manufacturer’s measurement and reflect the targeting location. Scale bars, 2 mm. **b**, The axial view of the mouse brain highlights the hippocampus, including both the dorsal and ventral regions. Overlapping the hippocampus are the FUS beams, depicted as circles representing the exact size of the beam.

The selected sonication parameters had a peak negative pressure of 0.63 MPa, a frequency of 1.5 MHz, and a 10 ms burst length with 30 burst period. These parameters corresponded to a mechanical index (**MI**) of 0.51, as determined by the transducer calibration provided by the manufacturer (FUS Instruments, Toronto, Canada). A total of two bolus MB injectiSons were conducted on each animal to cover each hemisphere, and we introduced a five-minute gap between each MB injection and insonation to allow the elimination of residual MB. To rapidly assess the feasibility and safety of the BBB opening parameters, we used FUS-BBBO with Evans blue dye (**EBD**) injection. EBD is an easily detectable dye frequently used to evaluate the efficiency of FUS-BBBO using histology [39]. We targeted FUS-BBBO with EBD in 4 male mice with a total of 96 opening sites and found that, after perfusion, 100% of the sites showed successful delivery of the EBD, suggesting that the chosen parameters were useful for reliable BBB opening with no gross tissue damage in any of the sites (**Supp Fig. 2a and 2b**, n = 2 male mice).

After validating FUS-BBBO parameters, we moved to administering the viral injections in a separate group of mice to investigate ATAC’s efficacy when coupled with our novel viral vector - AAV.FUS.3. We deliver the inhibitory DREADD receptor hM4Di (**DREADD-Gi**) fused to the fluorescent reporter mCherry, enabling histological visualization. Expression was driven by the calcium/calmodulin-dependent protein kinase II alpha (**CaMKIIa**) promoter to target excitatory pyramidal neurons selectively in the HPC. To assess the neuron inhibition ability of ATAC with AAV.FUS.3, we implemented the FUS targeting script to transduce DREADD-Gi throughout the whole HPC of the mouse and assessed the effect of CNO administration on the flurothyl-induced generalized seizure model. In particular, we measured the seizure threshold differences between ATAC-treated and untreated mice. Flurothyl is a volatile chemoconvulsant that acts as a GABAa antagonist, historically used to induce seizures in severely depressed patients as an alternative to electroconvulsive shock therapy [40], and it has been widely adopted in research as a valid ictogenesis model [41-43]. Since the HPC plays an essential role in seizure pathology [44], we hypothesized that inhibiting its excitatory pyramidal neurons non-invasively using the ATAC strategy would reduce the basal activity of those neurons, delaying the seizure onset.

To obtain ATAC mice, wildtype animals were immediately insonated with FUS accompanied by i.v. administration of AAV.FUS.3 containing DREADD-Gi-mCherry under the CamKIIa promoter (**Fig. 3a**). A group of wildtype mice (**Fig. 3b**) that received i.v. AAV injection without the exposure to FUS was included as a control (**Fig. 3a**). After two weeks, mice underwent the flurothyl seizure induction protocol. In this protocol, mice received AAV injection, including both exposed and not exposed to FUS (**AAV + CNO group**), received a single dose of 5 mg/kg CNO intraperitoneally (**i.p**.) sixty to seventy-five minutes prior to seizure induction to ensure a high plasma concentration of CNO [26, 45]; wild-type animals did not receive any injectable chemicals. After CNO incubation, the mouse was placed into the induction chamber (**Fig. 3c**) for at least one minute of habituation before the induction of seizures. The chamber is connected to a syringe pump that contains 10% of flurothyl ether in a glass syringe (**Supp Fig. 3a**). We recorded the time with videos of each individual mouse from the onset of the flurothyl enters the chamber (**Supp Fig. 3b**) until the animal showed a stage four (sitting) seizure severity according to the modified Racine scale [46]. The animal was then placed into a separate cage after they reached stage four of seizure severity for recovery and perfused after 90-120 minutes for histology examination. Two blinded reviewers reviewed the videos to ensure the subjectivity of the scoring. In this seizure model, wildtype mice have an average 358 second latency to stage four seizure. In contrast, ATAC mice have a significant increased latency with an average of 412.6 second (15.25% increase, P = 0.0018, **Fig. 3d**). Additional control showed that the activation of any DREADDs potentially expressed outside FUS-targeted brain regions (after a systemic AAV.FUS.3 injection, but in the absence of FUS-BBBO) did not result in changes to the seizure threshold (P = 0.98, **Fig. 3d**). This result confirmed the necessity of FUS in our ATAC treatment and validated the robustness of our flurothyl seizure induction protocol with n > 20 tested animals.

**Figure 3.**
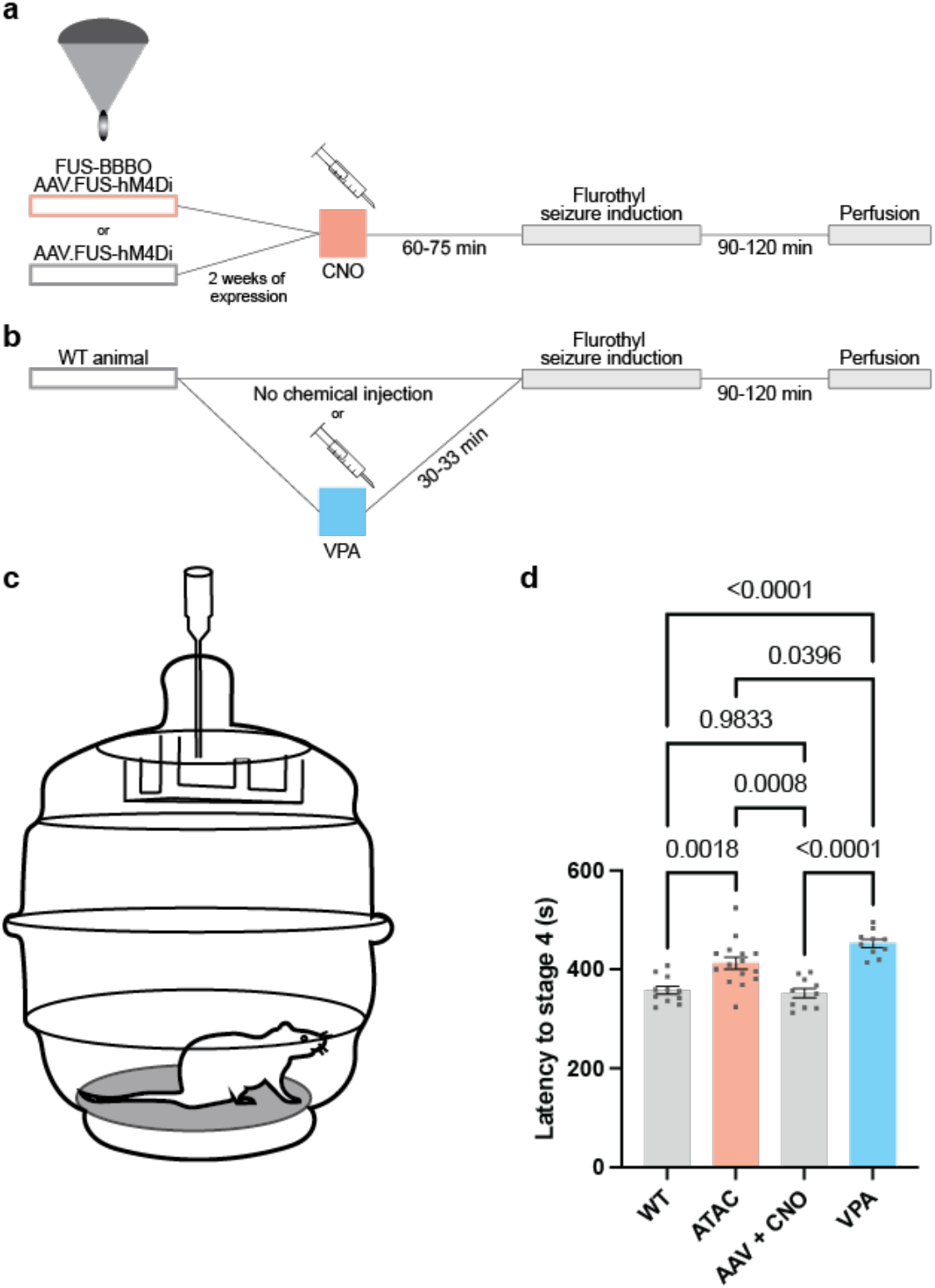
Flurothyl seizure threshold determination with ATAC and comparison with VPA. **a**, Two weeks after FUS-BBBO and the administration of AAV.FUS.3-DREADD (hM4Di-mCherry) mice were injected with CNO, then seizures were induced with flurothyl 60-75 minutes after CNO injection. Mice that received AAV.FUS.3 intravenously without FUS exposure underwent the same procedure. **b**, Wildtype animals that did not received any chemicals were also induced with flurothyl exposure. Mice that received VPA were induced thirty minutes after injection to match the T_max_ of this compound. All animals were cardiacally perfused between 90 - 120 minutes after the onset of the induction for histological analysis. **c**, Graphical representation of the flurothyl induction chamber. An 18G metal needle was inserted at the top of a transparent vacuum desiccator, a customized 3D-printed gauze holder was attached below the needle, and a new gauze was placed on the holder before each induction. **d**, Flurothyl generalized seizure latency to stage four of seizure severity. Groups are compared to each other using a Tukey honestly significant difference post hoc test (F (3, 42) = 19.04). No latency difference was found between WT mice and AAV + CNO mice (P = 0.9833). Significant increases in latency were found both in ATAC and VPA-treated mice compared with WT (P = 0.0018 and P < 0.0001, respectively) and the AAV+ CNO injected group (P = 0.0008 and P < 0.0001, respectively). An improvement was also found between ATAC and VPA-treated mice (P = 0.0396). The bar graph represents the mean ± s.e.m.

To benchmark our ATAC treatment with currently available AEDs to determine neuron inhibition efficacy, we chose VPA as our reference. VPA is a second-generation AED that is suitable for short-term and long-term usage in the treatment of generalized seizures and is often used as a benchmark to determine the efficacy of the third generation of AEDs [47]. Unlike ATAC, VPA acts throughout the brain, rather than only in the hippocampus, potentially contributing to its side effects [48]. We injected at a concentration of 130 mg/kg, which falls within the clinically prescribed dosage based on previously published mouse-to-human conversion [49, 50]. Mice that received the single i.p. injection of VPA underwent the same flurothyl seizure induction to stage four seizure severity based on the published protocol and the time to maximum response of VPA (Tmax) (**Fig. 3b**) [42, 51], ensuring the effective plasmid concentration. Mice treated with VPA showed an average seizure latency to stage four of 452.5 second, which has a 26.39% improvement compared with the wildtype animal (P < 0.0001, **Fig. 3c**). When compared with the efficacy of AAV.FUS.3 and ATAC, VPA-treated mice showed 9.66% better performance than ATAC-treated mice (P = 0.04, **Fig. 3c**). We did not observe any statistical differences in seizure latency across the blinded scorers who did not participate in any of the flurothyl induction experiments, including compound administrations (P > 0.9; **Supp Fig. S4a - 4d**). Moreover, across all four groups of animals, we did not observe any sex difference in seizure susceptibility (**Supp Fig. S5a-5d**).

After the behavioral assessment, all animals were perfused, and brains were extracted for histological examination. With immunostaining, the targeted locations, shown as an approximate FWHM beam size (**Fig. 4a**), had widespread hM4Di expression in histological sections, covering most hippocampal regions and a small segment of cortex above and thalamus below the dorsal hippocampus or next to the ventral hippocampus (**Fig. 4b**). Expression was especially strong in the molecular and polymorph layers of the dentate gyrus, the stratum oriens and stratum radiatum of the **C**ornu **A**mmonis 1 (**CA1**), CA2, and CA3 fields, as well as the pyramidal cells of the dentate gyrus (**DG**), CA1, CA2, and CA3 (**Fig. 4c**). In comparison, mice that received an i.v. injection of the same AAV.FUS.3 vector and CNO but without FUS-BBBO showed no detectable fluorescent signal in these brain regions, confirming that BBBO was required for gene delivery at the viral dose used in our study (**Fig. 4d**).

**Figure 4.**
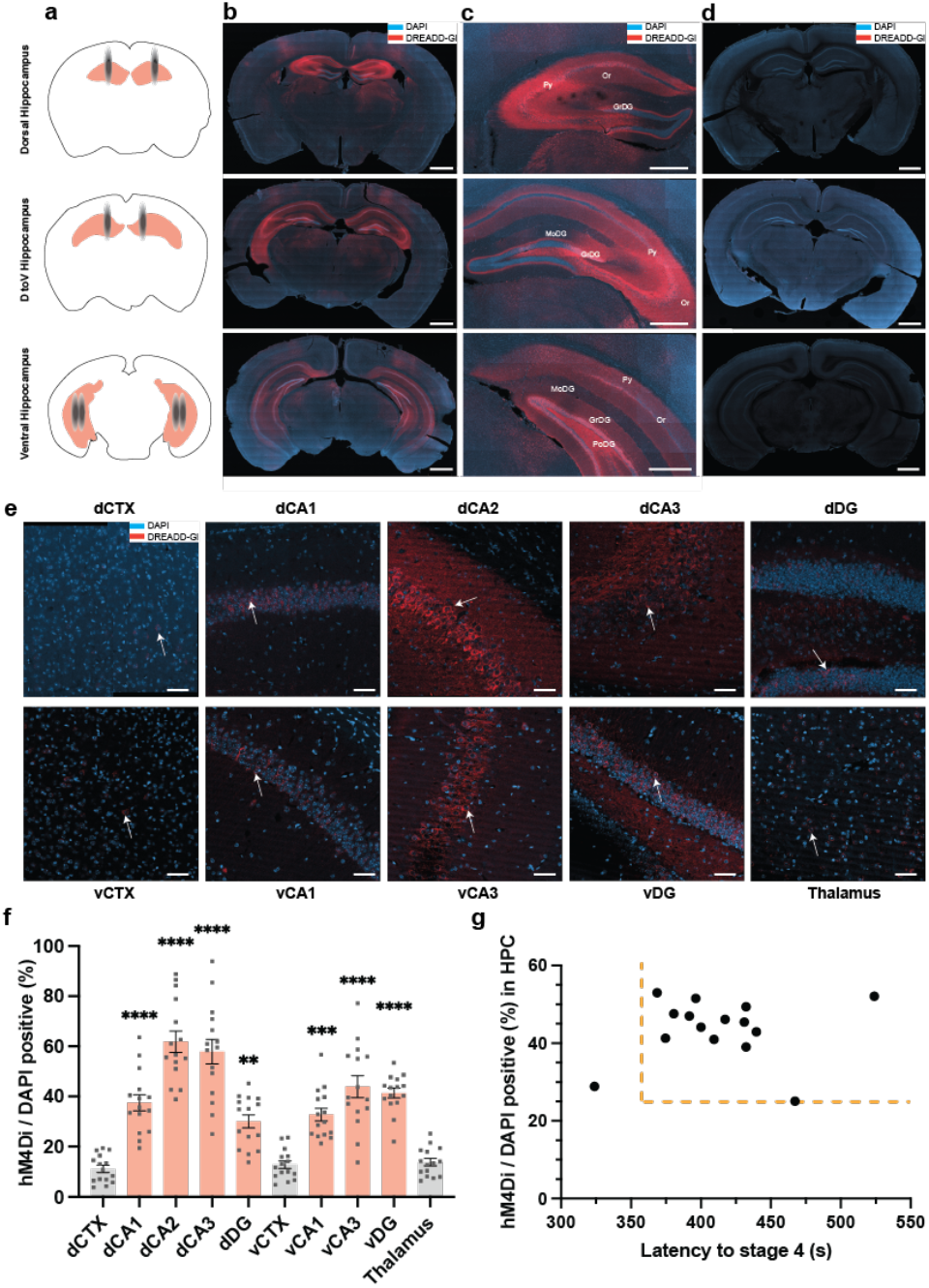
Spatial specificity and efficacy in neuronal modulation of hM4Di expression. a, Graphical representation of FUS beams with an indication of the FWHM pressure of the beam in a grey scale, where a region with a darker color represents a higher peak negative pressure. b, Representative brain sections from two animals showing dorsal (top), dorsal to ventral transition (middle), and ventral (bottom) hippocampus immunostained for hM4Di-mCherry (red) two weeks after FUS-BBBO and injection of AAV.FUS.3 encoding this hM4Di-mCherry under the CaMKIIa promoter. The DAPI stain demarcates cell nuclei (blue). Scale bar, 1 mm. c, Magnified view of dorsal (top), dorsal to ventral transition (middle), and ventral (bottom) hippocampus showing widespread expression in oriens (Or), pyramidal (Py), granule (GrDG), polymorph (PoDG), and molecular (MoDG) cell layers of the hippocampus. Representative of n = 3 independent brains analyzed. Scale bars, 500 μm. d, Representative immunostaining result for hM4Di-mCherry in a mouse that received the same viral construct and CNO injection, but did not undergo FUS-BBBO. Representative of n = 2 independent brains analyzed. Scale bar, 1 mm. e, Representative images of mCherry fluorescence (red) in each field. The DAPI stain (cyan) marks cell nuclei. Arrows show examples of DREADD-positive cells. Representative of n = 2 mice. Arrows indicate the DREADD-positive cell bodies. Scale bars, 50 μm. f, Percentage of cell bodies with detectable mCherry fluorescence in pyramidal layers of the hippocampus and overlaying FUS-targeted dCA1 and vCA1, and thalamus as an untargeted negative control. These values indicate relative transduction efficiency in different fields of the hippocampus. N = 15 mice; one-way analysis of variance test compared with thalamus, variance ratio F (9, 140) = 33.92; **P < 0.01, ***P < 0.001, ****P < 0.0001 in comparison with negative control (thalamus), using a Dunnett honestly significant difference post hoc test. The bar graph represents the mean ± s.e.m. g, A correlation graph that describes the relationship between DREADD expression in the whole HPC region and seizure latency. The vertical dash line is on the 358s mark on the x-axis, which is the average latency of WT mice; the horizontal dash line is on the 25.07% mark of the y-axis, which is the minimal whole hippocampus transduction rate that produces the seizure latency improvement effect calculated by averaging the transduction rate of CA1-3 and DG of both dorsal and ventral HPC.

A quantitative comparison of expression in FUS-targeted areas across all 15 ATAC-treated mice was performed using mCherry fluorescence in cell bodies of granular and pyramidal cell layers, which allowed for a direct comparison of transfection efficiency between hippocampal regions. Our analysis showed that, on average, more than 50% of the cells in dorsal CA2 (**dCA2**) and CA3 were successfully transfected, and that dorsal and ventral dentate gyrus contained 30% and 41% positive cell bodies, respectively (**Fig. 4e, 4f**). The cortex had lower transfection efficiencies, suggesting that the region is less susceptible to transfection after BBBO than other regions, or that simply the lower-pressure areas of the FUS beam, away from the center of targeting, were less efficient at AAV delivery. As a representative non-targeted region, we looked for expression in the thalamus, which was shown in previous studies to be particularly susceptible to transfection following systemic delivery of AAV9 [52], and found similar positive cell counts compared with both dorsal and ventral cortex (% positive = 13.8%, P > 0.99, **Fig. 4f**), indicating the precise spatial targeting of our FUS-BBBO protocol. This expression pattern is expected, as the center of the ultrasound beam corresponds to a peak negative pressure (**PNP**) of 0.63 MPa, which gradually decreases toward the edges, reaching approximately 0.315 MPa at FWHM distance, resulting in lower transduction rates at the beam periphery, such as the thalamus. Full results of the statistical tests can be found in **Supplementary Table 1**. Next, we drew a distribution graph showing the correlation between the whole HPC transduction efficiency and stage four latency. We used the average latency of wildtype animals (358 seconds) as the neuronal inhibition threshold. We found that the minimal transduction rate of the entire HPC required for AAV.FUS.3 to produce an inhibitory effect was 25.07%, which is the average transduction rate of dorsal and ventral CA1-3 and DG. Notably, 93.33% (14 out of 15) of animals with transduction rates greater than 25.07% exhibited longer latencies to reach stage four compared to wildtype animals, indicating an effective neuronal inhibition (**Fig. 4g**).

To evaluate the safety and adverse effects of ATAC and whole HPC targeting with AAV.FUS.3, we examined the weight loss of wildtype, ATAC, and AAV + CNO mice. Any significant loss of body weight would serve as an indication of substantial adverse effects of ATAC on the well-being of animals, with 15% of the weight loss consisting of a humane end-point (**Fig. 5a**). After the FUS insonation and AAV injection, we monitored the weight of the animals for four consecutive days and at the day of the experiment. For wild-type animals, we also monitored body weight throughout the experiment over the same timeframe. No significant weight loss was observed in any group, and body weight increased over time and remained above the humane endpoint threshold (Fig. 5a). We further analyzed the safety profile with neutral buffered formalin fixed tissues. We initially examined all sections from ATAC-treated mice using a fluorescence microscope. We then selected the sections with the brightest mCherry fluorescence signals and within 200 μm from that section for further analysis, which indicated the location of highest FUS pressure and thus likely the center of the beam. We reasoned that fluorescence intensity correlates with the efficiency of FUS-BBBO, as increased fluorescence suggests greater AAV.FUS.3 penetration through the BBB. Consequently, these regions, may also be more susceptible to potential damage caused by either FUS or AAV.FUS.3. For mice that received AAV but were not exposed to FUS, sections were selected based on anatomical structure. We first performed hematoxylin staining, and found no visible tissue damage differences between ATAC mice and AAV + CNO mice, demonstrating the safety of our FUS-BBBO perimeters in our study (**Fig. 5b and 5c**). We then evaluated the potential inflammation and neuronal damage by staining with GFAP (astrocyte marker), IbaI (microglial marker), and NeuN (neuronal marker). To quantify the GFAP and IbaI expression, we used positive pixel analysis to focus on the CA2 region of both hemispheres and combined the counts from both hemispheres, as the complex morphology of glia and microglia made cell counting more complicated. GFAP-positive pixel analysis showed a significant 2.77±0.34-fold (SEM) increase of astrocyte activation in ATAC-treated mice but not in the AAV + CNO mice (P < 0.0001, **Fig. 5e and 5g**). When comparing the glia expression between the same group, we noticed that the expression of the IbaI marker is a 1.73±0.3-fold (SEM) increase in AAV + CNO group (P = 0.025, Fig. 5e and 5h). NeuN quantification was done by two blinded experimenters who did not participate in the behavior experiment. We quantified the number of neurons in the pyramidal layer of CA2 regions by manual counting (**Fig. 5e**), and we didn’t notice statistically significant differences between the groups (P = 0.1236, **Fig. 5i**), indicating that AAV.FUS.3 or FUS-BBBO did not generate any neuronal damage during the treatment.

**Figure 5.**
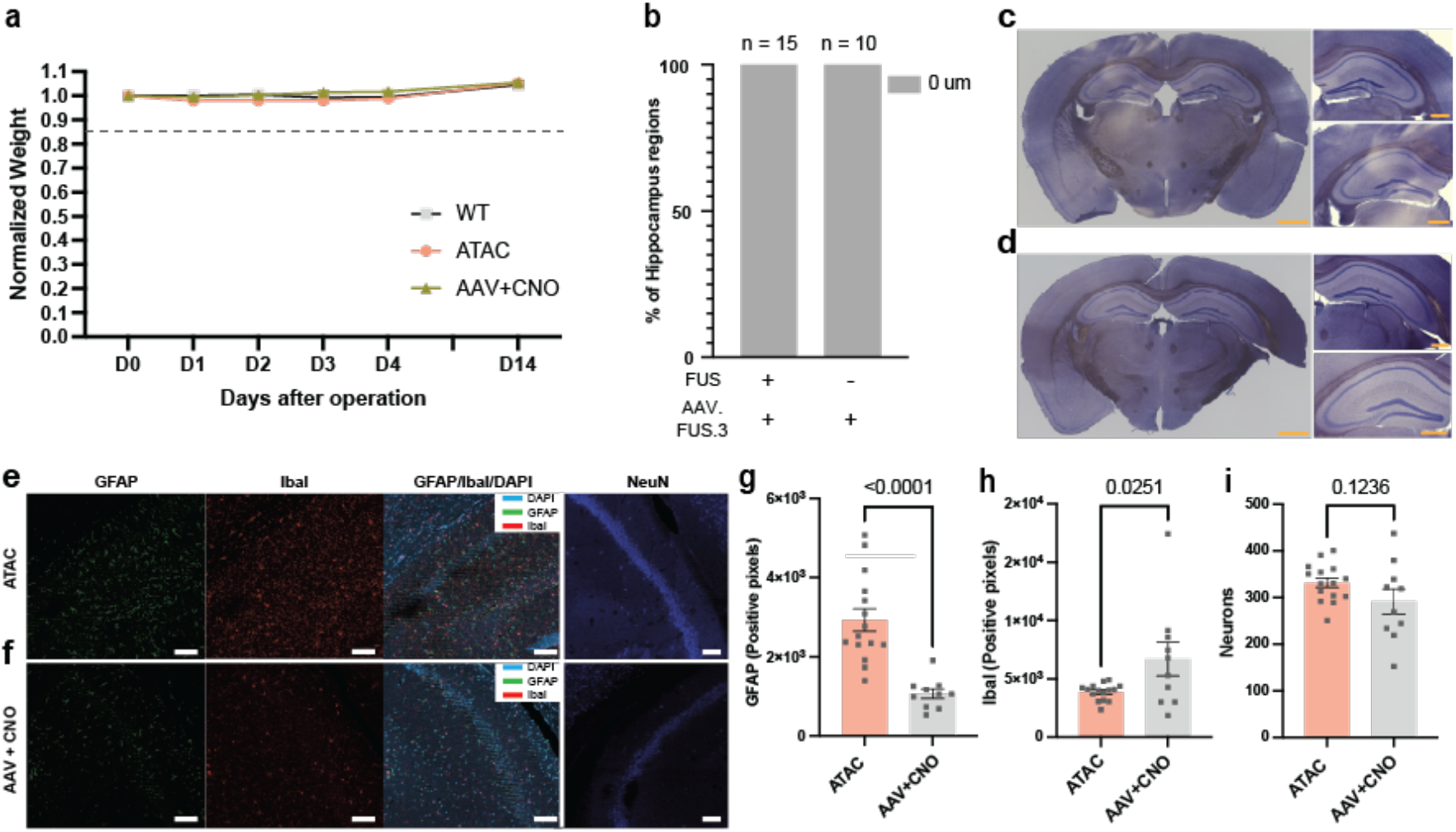
Long-term safety profile of ATAC and AAV.FUS.3. **a**, All mice were monitored for four consecutive days after surgery, and on the day of seizure induction after they were insonated with FUS or only received a viral injection for any weight loss or abnormal activity. No significant weight loss was observed. The dashed line indicates the humane endpoint due to weight loss. **b**, Histological classification of brain tissue damage of FUS targeted sites in 15 ATAC-treated mice, and 10 mice received the AAV injection without FUS exposure. **c**, Representative hematoxylin-stained brain section of ATAC-treated mice. The left image is the whole brain image, and the images on the right are ROIs of both insonated hippocampus regions. Scale bar, 1000 μm for the left image and 500 μm for the right ROIs. **d**, Representative hematoxylin-stained brain section of mice received AAV and CNO but were not exposed to FUS. Scale bar, 1000 μm for the left image and 500 μm for the right ROIs. **e**, Representative 20x confocal images of the hippocampus CA2 region showing the ROI for GFAP (astrocyte marker, in green) and IbaI (microglia, in red) analysis for ATAC-treated mice after immunostaining. The DAPI stain (cyan) marks cell nuclei. Images for NeuN (neuronal marker, blue) were taken separately with different sections. Scale bar, 100 μm. **f**, Representative 20x confocal images of the hippocampus CA2 region of mice received AAV and CNO but were not exposed to FUS with the same types of immunostaining as the ATAC-treated group. **g**, GFAP was analyzed using a positive pixel count with ImageJ in all groups. Groups are compared using a two-tailed heteroscedastic t-test, showing an increased GFAP expression in ATAC-treated mice compared with the IV-infused mice (P < 0.0001). **h**, IbaI was analyzed using a positive pixel count with ImageJ in all groups. Groups are compared using a two-tailed heteroscedastic t-test, showing a small increase in IbaI expression in mice receiving AAV and CNO but not FUS exposure compared with the ATAC-treated mice (P = 0.0251). **i**, Number of neurons in the CA2 pyramidal layers of both hemispheres. Groups are compared using a two-tailed heteroscedastic t-test, showing no statistical difference between ATAC-treated and mice that received AAV and CNO without FUS exposure (P = 0.1236). All bar graphs represent the mean ± s.e.m.

## DISCUSSION

Taken together, our results establish ATAC, in combination with AAV.FUS.3 offers a unique combination of spatial, cell-type, and temporal specificity without tissue damage or neuronal loss. When used in as a tool to modify seizure threshold, the hippocampus-specific treatment was comparable, albeit slightly less effective, to that of broadly-acting valproic acid administered at clinically prescribed doses in the same seizure model.

In contrast to traditional intraparenchymal viral delivery, FUS-BBBO allows comprehensive transduction of the entire hippocampus in just two sessions with minimal tissue damage and high scalability to larger species [53]. Invasive intraparenchymal delivery, on the other hand, often requires multiple brain penetrations to adequately target the unilateral hippocampus with preferred tropism [54], even when using convection-enhanced delivery in large animals [32, 55]. Furthermore, FUS-mediated delivery reduces the risk of local toxicity associated with high viral particle concentrations near injection sites, thereby minimizing potential side effects. Compared with emerging ultrasonic neuromodulation techniques, which rely on direct ultrasound stimulation of brain regions or ultrasound-mediated drug delivery, ATAC offers several key advantages. It does not require animals to remain stationary under a mounted ultrasound transducer during modulation [56-58]. Instead, following transduction and expression of chemogenetic receptors in a genetically defined subset of cells at the FUS-targeted site, neuromodulation can be conveniently achieved through systemic administration of a bioavailable ligand. In comparison to optogenetics and deep-brain stimulation, which also provide spatial and temporal control, ATAC enables non-invasive neuromodulation without the need for a chronically implanted device such as an optical fiber, which significantly reduces invasiveness and simplifies long-term applications.

Overall, given that AAV-based therapies are currently under investigation in numerous clinical trials [59-61], FUS-BBBO is actively being implemented in clinical settings [13, 62], and DREADD efficacy has been validated in non-human primates [32, 63], the ATAC approach using AAV.FUS.3 holds a clinical promise.

In our behavioral proof of concept, a single injection of CNO two weeks after the FUS-BBBO procedure resulted in a 15.25% improvement in the flurothyl seizure threshold, and the dose of CNO did not contribute to the change in seizure threshold, validating the efficacy of neuromodulation of AAV.FUS.3. When we benchmarked the efficacy of VPA with the same protocol, ATAC was 9.66% less potent than VPA. We hypothesize that the limited seizure attenuation observed in some animals may be due to seizure initiation in brain regions not targeted by ATAC, such as the amygdala or cortex. In contrast, systemic administration of small-molecule drugs affects the entire brain. We selected VPA as our benchmark because of its broad-spectrum antiepileptic activity and well-documented anti-epileptogenic effects across various seizure models, including pentylenetetrazole (**PTZ**), pilocarpine, kainic acid, amygdala kindling, and, most importantly, in the flurothyl model [42, 47, 64]. Since both PTZ and flurothyl are GABAa receptor antagonists [57, 65], we reasoned that the mechanism of epileptogenesis is relatively similar. The seizure-suppressive efficacy achieved with a similar dose of VPA in our model aligns with results from previously published studies [42], supporting the robustness and reliability of our experimental system. Notably, the dosage of antiepileptic drugs, such as VPA, prescribed for effective chronic seizure management is often associated with undesirable side effects, including both neurological and liver toxicity [48]. In contrast, DREADDs require a very low concentration of a small-molecule ligand to activate the receptor and exert a suppressive effect. Although CNO can be back-metabolized into clozapine, adverse effects are typically observed only at high doses, around 100 mg per day in humans [66] or 20 mg per kg in mice, significantly higher than the dose required for effective DREADD activation. With postmortem analysis, we concluded that with a 25.07% pyramidal neuron transduction rate, there is a 93% chance that our therapy will be effective **(Fig. 4g)**. We did not exclude any animals in our study, even though there is one animal that has less than 30% dorsal CA3 transduction rate, which is the effective threshold for modulation [67], because we targeted the entire hippocampus, and the region of interest (**ROI**) we chose are random and may have missed the highly transduced region.

We also observed no significant weight loss in ATAC or AAV + CNO mice compared with the wildtype animals (**Fig. 5i**), indicating the FUS-BBBO procedure and AAV.FUS.3 vector were well tolerated and did not result in overt systemic side effects. However, we found an increase in GFAP expression in the CA2 region of ATAC-treated mice compared to AAV + CNO mice (**Fig. 5g**). We attribute this to a potential immune response triggered by the immunogenic nature of the AAV.FUS.3 vector due to foreign protein expression [68], and possibly amplified by the mechanical effects of FUS at higher pressures [69], but not with the expression of DREADD, as other papers reported the GFAP immunoreactive region is comparable in DREADD-expressing and non-expressing animals [70]. However, such an immune response is transient [71] and can be alleviated with anti-inflammatory corticosteroid injection [72]. Additionally, we observed elevated Iba1 expression in the AAV + CNO group, without FUS-BBBO (**Fig. 5h**), which may reflect individual variability or inflammation related to seizure induction rather than the delivery method, as other researchers showed that the immune response time of microglia is faster than that of astrocytes in the seizure model [73]. Importantly, no neuronal loss was observed in any group treated with ATAC and AAV.FUS.3, further reinforcing the safety of this approach for non-invasive neuromodulation.

The ATAC with AAV.FUS.3 paradigm could be further enhanced through improvements in its core components: FUS-BBBO procedure, AAV vectors, chemogenetic receptors, and ligands. For instance, the safety and efficiency of FUS-BBBO can be improved by integrating real-time feedback control based on subharmonic emissions of microbubble oscillation, which helps regulate peak negative pressure [74, 75]. Additionally, smaller contrast agents, such as nanobubbles, enable more homogeneous BBB opening by penetrating smaller vessels [74]. Further optimization of AAV vectors is also needed to improve transduction efficiency and reduce the required viral dose. This includes developing compact, cell-type-specific promoters that remain robust across species, particularly in non-human primates, to facilitate the translational properties [76, 77]. In parallel, ongoing studies are refining chemogenetic receptors and their binding ligands. For example, new DREADD ligands, such as deschloroclozapine [75] could reduce potential side effects, and newly engineered receptors responsive to ligands such as salvinorin B [78] could allow for multiplexed modulation

These innovations will significantly enhance the versatility and clinical translatability of ATAC and AAV.FUS.3 as a non-invasive, precise method for neuromodulation. However, the efficacy of AAV.FUS.3 in chronic disease models remains unevaluated, and optimal dosing strategies must be established to account for potential synaptic desensitization over time.

## MATERIALS AND METHODS

### Animals

Fifty wild-type C57BL/6J mice (31 male and 19 female) were used in this study. Mice were purchased from Jackson Laboratory. Animals were housed in a 12 h light/dark cycle with lights on at 7:00 AM and were provided with water and food ad libitum. Animal experiments were conducted under a protocol (#IACUC-23-265-RU), in accordance with NIH guidelines, and approved by the Institutional Animal Care and Use Committee of Rice University.

### AAV production

AAV.FUS.3 vector was packaged by adopting a published protocol [79]. In brief, HEK293T cells (AmericanType Culture Collection, CRL-3216) were transfected with the DREADD plasmid containing inverted terminal repeats of AAV (pAAV): pAAV-CaMKIIa-hM4Di-mCherry (Addgene plasmid # 50477), AAV.FUS.3 (Addgene plasmid # 222333) and pHelper plasmids. After 24 hours, cells were exchanged with a fresh DMEM (Corning, 10-013-CV) supplemented with 5% FBS and non-essential amino acids (Gibco, 11140050). At 4 days after transfection, cells were harvested, and media were mixed with 1/5 volume of PEG solution (40% PEG 8,000, 2.5M NaCl) to precipitate AAV at 4 °C for 6 hours. Cells were resuspended in PBS and lysed by the freeze-thaw method. The precipitated AAV was pelleted by centrifugation, resuspended in PBS, and combined with these cells. The combined lysate was added with 50 U/mL Benzonase (Sigma-Aldrich, E1014) and incubated at 37 °C for 45 min. AAV purification was carried out using iodixanol gradient ultracentrifugation. A Quick-Seal tube (Beckman Coulter, 344326) was loaded with the iodixanol gradients (Sigma-Aldrich, D1556), including 60%, 40%, 25%, and 15%. The lysate was centrifuged, and the resulting clarified lysate was loaded on top of the iodixanol layers. The tube was sealed and centrifuged at 350,000g for 2.5 hours using a Type 70 Ti fixed-angle rotor of an ultracentrifuge (Beckman Coulter). AAV was collected by extracting the 40– 60% iodixanol interface and washed using the Amicon centrifugal filter unit with a 100-kDa cutoff (Sigma-Aldrich, UFC910024). The final AAV was filtered by passing through the 0.22-μm PES membrane. Viral titers were determined using the qPCR method.

### FUS-BBBO

C57BL/6J 16–18-week-old, male and female, mice were anesthetized with 2-3% isoflurane in the air. The fur on their heads was removed using a trimmer. Next, a premade catheter with a 30-gauge needle washed with 10U/mL heparin and 0.9% saline was placed in the lateral tail vein and secured with tissue glue. Subsequently, the mice were placed on a stereotactic instrument, and their heads were fixed with ear bars. The incision area was washed thrice with a chlorhexidine scrub, followed by a chlorhexidine solution. The mice were injected via the tail vein with Evans blue dye (Sigma-Aldrich, E1219, 0.2 mg per g) or AAV.FUS.3 (1E10 viral particles per g) encoding DREADD-Gi in the plasmids. Immediately after intravenous injection of the AAV, mice were injected with approximately 0.72 × 10^6 Definity (Perflutren Lipid Micro-sphere, Lantheus Medical Imaging, Billerica, MA, USA) microbubbles (MB) per gram of body weight. Definity was dissolved in sterile saline (20 μL of MB in 980 μL of saline solution, Hospira, 00409-4888-10) [12] to allow for the administration of more manageable volumes. The transducer, coupled to the head via Aquasonic gel (Parker labs, Fairfield, NJ), was placed on the top of the skull of the animal. The targeting depth was adjusted using the physical movement of the transducer through a software interface to target specific brain regions. The transducer used in the experiment had an FWHM pressure of ∼1.0 × 4.9 mm and an approximate volume of 2.56 mm3.

The ultrasound parameters included a 1.5 MHz stimulation frequency and a peak-negative pressure of 0.63 MPa. The provided pressures were based on preliminary experiments in our lab, validated with EBD results, and were not derated for the skull attenuation, as the exact pressure at the brain will vary depending on the location. Each site was insonated 30 times with a 10 ms burst duration. Using custom software, we enabled the (RK-50 FUS Instruments, Toronto, Canada) to target each half of the hippocampus region in a single session in a relatively symmetrical manner. Bilateral hippocampi were targeted separately, and each insonation session was designed to finish within 2 min to achieve the highest efficacy of MB for BBBO [80]. A five-minute gap was introduced between the first and the second insonation sessions to avoid the inconsistency of BBB opening efficiency due to the accumulation of remaining MB in the circulation in the previous injection. Each site was insonated for 10 ms, and then the transducer moved to the next site until it reached the last site. The insonation cycle was repeated 30 times. Due to the limitations of our instrument, the transit time of the transducer between different brain sites varied due to the differences in size and shape of the insonated brain region, as our sites were input manually to show a symmetrical opening on HPC bilaterally. A total of 12 sites per hemisphere script were used on all the FUS-treated animals, and microbubbles were injected per hemisphere per insonation. The exact paths and durations of insonation procedures are shown in **Figs. 2a and 2b**, and the insonation order is shown in **Supp Fig S1**. The EBD results are shown in **Supp Fig S2**. The mice were monitored, and their body weights were recorded for four days following surgery and then on day 14 before flurothyl seizure induction.

### Drug administration

Water-soluble CNO (Hello Bio, HB6149) and valproic acid (Sigma-Aldrich, 676380) were each dissolved in saline (Hospira, 00409-4888-10) to prepare working solutions on the day of injection. CNO was dissolved at 1 mg per mL, while VPA was dissolved at 20 mg per mL. Both solutions were filtered through a 0.22 μm filter for sterilization before injection.

### Flurothyl Seizure threshold determination

Mice that underwent the full ATAC treatment, as well as those that were not exposed to FUS but instead received a viral vector, were injected with CNO (5 mg per kg, i.p.) and allowed to incubate for 60-75 minutes to allow CNO to reach its pharmacokinetic peak [45]. For mice treated with VPA, VPA was administered (130 mg per kg, i.p.) with a 30-33 minute incubation time before flurothyl seizure induction. This specific timeline was selected to ensure CNO and VPA reached their respective pharmacokinetic peaks before seizure induction.

The seizure induction time point was chosen based on previous studies about AAV.FUS.3 expression [36]. Two weeks after ATAC or IV viral injection, individual mice (16-20 weeks old; WT: 17-20 weeks, ATAC: 18-20 weeks, AAV + CNO: 17 weeks, and VPA: 16 weeks) were exposed to 10% flurothyl (bis(2,2,2-trifluoroethyl) ether, Sigma-Aldrich, 287571). A 10% flurothyl solution was made by diluting flurothyl ether in 95% ethanol (KOPTEC, V1101) based on a published protocol to make a 50 mL flurothyl solution and stored in an air-tight glass bottle for future usage [65]. Flurothyl seizure was induced by adopting the same protocol but with modification (**Supp Fig. S3**). Mice were transferred to the experimental room 1 h before the experiment. Mice were placed in the induction chamber 60-85 seconds before induction for habituation. Flurothyl was filled in a glass syringe and pumped via a syringe pump at a speed of 7.8 mL per hour into a small all-clear vacuum desiccator with an estimated volume of 14.4 liters (Ted Pella, 2240.1). Petroleum jelly was applied on the top and the bottom of the chamber to ensure an adequate seal. A customized 3D printed gauze holder was glued at the top of the lid to carry a gauze for the flurothyl liquid to evaporate. One mouse at a time was induced in the flurothyl chamber using a new gauze pad for each induction. A modified Racine scale was employed for seizure threshold determination, with the following scoring criteria: stage 1, sudden behavior arrest; stage 2, facial jerking; stage 3, neck jerks; stage 4, clonic seizure or sitting; stage 5, tonic-clonic seizures (lying on the belly); stage 6, tonic-clonic seizure (lying on the side) or wild jumping; and stage 7, tonic extension, potentially leading to respiratory arrest and death [46]. The latency to stage 4 was measured from flurothyl liquid entering the chamber, and the mouse was placed into a separate recovery cage until consciousness was regained and the mouse was able to move freely. All flurothyl experiments were video recorded for behavioral assessment by two blinded reviewers and confirmed without statistical significance (S.L. and S.S.L., P > 0.892, **Supp Fig. S4a-4d**).

### Immunohistochemistry for evaluation of DREADD expression, BBBO efficacy, and immune response

After 90-120 minutes of seizure induction, mice were perfused with 10% formaldehyde neutral buffer (Sigma-Aldrich, HT501128) for 10 minutes. The brains were then extracted and fixed in 10% formaldehyde for 16 hours at 4 °C. Following fixation, brains were sectioned in the coronal plane using a vibratome (Leica VT1200S), with 50 μm serial sections obtained for further analysis. To visualize DREADD expression across brain regions, 50 μm tissue sections were treated with blocking solution (0.2% Triton X-100, 10% donkey serum in 1X phosphate-buffered saline (**PBS**)) for one hour, then incubated overnight at 4°C with polyclonal rabbit anti-mCherry antibody (ThermoFisher, PA534974, 1:1000) in blocking solution. After three 10-minute washes in 1X PBS, sections were incubated with donkey anti-rabbit secondary antibody conjugated to Alexa Fluor 647 (ThermoFisher, SA5-10041, 1:1000) for 3 hours at room temperature, washed again, and mounted on glass slides using Antifade Mounting Medium with DAPI (Vector Laboratories, H-1800). Imaging was performed using a Keyence BZ-X810 fluorescence microscope (Osaka, Japan) in the Cy5 channel under 20x magnification with sectioning mode, and representative images were taken with a confocal microscope (Zeiss LM 800). Images were acquired with consistent exposure and camera gain, and a uniform threshold was applied to each, ensuring standardized analysis across all images. Mice that received EBD injections were perfused 20 minutes later and fixed with the same procedure, but sectioned with 100 μm serial sections to better visualize the EBD color. Imaging was performed using a Keyence BZ-X810 fluorescence microscope (Osaka, Japan) under bright field and Tx Red channel with high resolution mode with 4x magnification.

For GFAP, IbaI, and NeuN staining, sections were selected from the ATAC and AAV + CNO groups. The ATAC-treated group was assessed under a fluorescence microscope, and the section with the highest transduced region was chosen, while sections from the AAV + CNO group were selected based on anatomical correspondence to the ATAC-treated group. Sections were treated with blocking solution (0.2% Triton X-100, 10% goat serum in 1X PBS) for two hours, then incubated overnight at 4°C with anti-GFAP (Abcam, ab7260, 1:500) and anti-IbaI (Abcam, ab283346, 1:1000) in blocking solution. Following three 10-minute washes in 1X PBS, sections were incubated with goat anti-rat Alexa Fluor 647 (ThermoFisher, A-21247, 1:500) and goat anti-rabbit Alexa Fluor 488 (ThermoFisher, A-11008, 1:500) for 2 hours at room temperature for GFAP and IbaI staining. For NeuN staining, sections were blocked identically and stained with conjugated anti-NeuN Alexa Fluor 405-conjugated antibody (Novus Biological, NBP1-92693AF405, 1:500) overnight at 4°C. After an additional series of washes, sections were mounted using Anti-fade Mounting Medium with DAPI (Vector Laboratories, H-1800) for the GFAP and IbaI sections. Sections stained with NeuN were mounted with Antifade Mounting Medium (Vector Laboratories, H-1700), and imaged using a Keyence BZ-X810 fluorescence microscope under 20x magnification with sectioning mode, and representative images and NeuN quantification images were taken with a confocal microscope (Zeiss LM 800) under 20x magnification. Images were acquired with consistent exposure and camera gain, and a uniform threshold was applied to each, ensuring standardized analysis across all images.

### Histological analysis

Hematoxylin staining was performed to evaluate the safety profile of ATAC. Sections (ATAC-treated n = 15 and AAV + CNO n = 10 mice) for hematoxylin staining were picked within 200 μm from the sections used for GFAP and IbaI staining, and sections from mice that received AAV and CNO without FUS exposure were chosen based on the anatomical location of ATAC mice.

Hemotoxicity staining was performed according to the following procedure. The tissue section was immersed in 100% ethanol for 1 minute, then transferred to 95% ethanol for 1 minute. A subsequent 1-minute rinse in distilled water was carried out. The section was then incubated in 50% hematoxylin (H&E Staining Kit, Abcam, ab245880) for 1 minute, after which it was rehydrated with a 3-minute rinse in distilled water. The tissue was then soaked in Bluing Reagent (H&E Staining Kit, Abcam, ab245880) for 1 minute, followed by a final 3-minute rinse in distilled water. The stained sections were mounted onto glass slides using Antifade Mounting Medium (Vector Laboratories, H-1700) and air-dried overnight in a dark environment before imaging. Imaging was performed using a Keyence BZ-X810 fluorescence microscope under 4x for the whole brain and 20x magnification for the hippocampus region with high resolution mode.

For a quantitative comparison of expression levels between various hippocampus regions, we used mCherry fluorescence localized to cytoplasmic compartments and counted the number of cells in the pyramidal layers of the hippocampus that showed detectable fluorescence. Cells that showed mCherry fluorescence surrounding the nucleus for at least 50% of its circumference were counted as positive to allow for a consistent comparison of expression between different hippocampal regions and conditions. The non-cytoplasmic localization of DREADD-mCherry necessitated the selection of this threshold. All the images were normalized in the background for a comparable expression evaluation. Expression was evaluated for randomly picked 3–5 sections per animal, and cells from each subfield of the hippocampus, cortex, and thalamus were added for each animal and normalized by the number of DAPI-stained cells in the granular cell layer of that subfield. ROI of regions being analyzed was the relatively hot spot in the hippocampus region, and CTX, and thalamus were right above or below the hippocampus as also insonated by FUS beams. The interexperimenter variability was determined for two different researchers (H.L. and S.N.) for n = 5 samples, with the difference in means smaller than 5% (mean = 31.7% versus mean = 36.1%, P = 0.07, heteroscedastic two-tailed t-test).

For GFAP and IbaI analysis, the bilateral dorsal CA2 region was selected as the ROI. ImageJ software was used to quantify total positive pixel counts across targeted sites with the same threshold within GFAP and IbaI analysis. For quantifying NeuN in the CA2 region, two blinded observers (S.N. and E.K.R.) manually counted the total number of neurons in a 2 × 2 image taken by 20x confocal of each animal in ATAC-treated mice and AAV + CNO mice. The backgrounds were maintained uniformly for accurate quantification. The CA2 region of both hemispheres in a single mouse was analyzed, and the total number of neurons was combined.

### Statistical analysis

Statistical analysis was performed using Prism (GraphPad Software, version 10.4.1). A two-tailed t-test with unequal variance was used to compare two datasets, while one-way ANOVA with Tukey and Dunnett’s honestly significant difference post hoc test was applied for comparisons involving more than two datasets. Statistical significance was set at *P < 0.05; **P < 0.01; ***P < 0.001; and ****P < 0.0001, unless stated otherwise. All P values and statistical test results are available in the Source Data. All data were tested for normality using a Shapiro–Wilk test. Figures were constructed using Adobe Illustrator. The MRI images in **Fig. 2a** are acquired from the Allen Common Coordinate Framework (**CCF**) and published data [81].

## Data availability

The authors declare that all data supporting the results in this study are available within the paper, its Supplementary Information, “Supplementary Figures S1-S5 and Table”. Microscopy images are available from the corresponding author upon reasonable request owing to their large size and number.

## ACKNOWLEDGEMENTS

The work was supported by the G. Harold & Leila Y. Mathers Foundation grant to J.O.S. We thank Ben Avants (Rice University) for the help with 3D printing and Dr. Zhimin Huang (University of Pittsburgh) for the helpful discussion on FUS parameter design.

## AUTHOR CONTRIBUTIONS

J.O.S. and H.L. conceived and planned the research. H.L. and J.O.S. designed the experiments and wrote the paper, with input from all other authors. H.L. performed and participated in all experiments described in the study. S.L. and S.S.L. viewed the videos and recorded the seizure latency. S.N. and E.K.R. analyzed histological images.

