## Supplementary information for "Noninvasive Control of Seizure Threshold with Acoustically Targeted Chemogenetics"

### Supporting Information

**Supplementary Fig. S1**

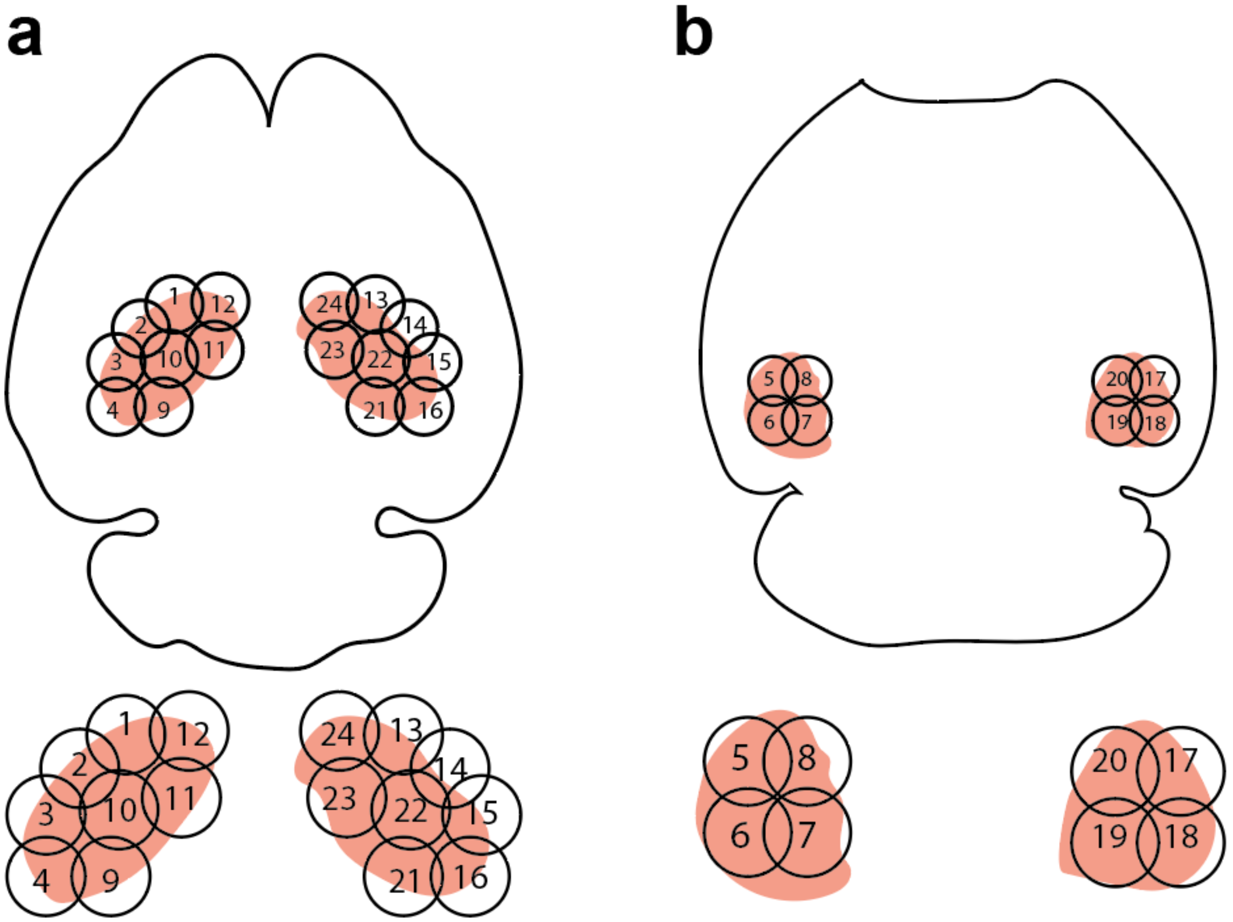


**Supplementary Fig. S1. Targeting strategy for whole hippocampus AAV.FUS.3 delivery.** The axial view of the mouse brain highlights the hippocampus. Each beam is labeled with a number, indicating the insonation order ranging from 1 to 12. The contralateral hemisphere was insonated with the order from 13 to 24. a. Axial view of dorsal hippocampus overlapped with FUS beams. b. Axial view of the ventral hippocampus overlapped with FUS beams.

**Supplementary Fig. S2.**

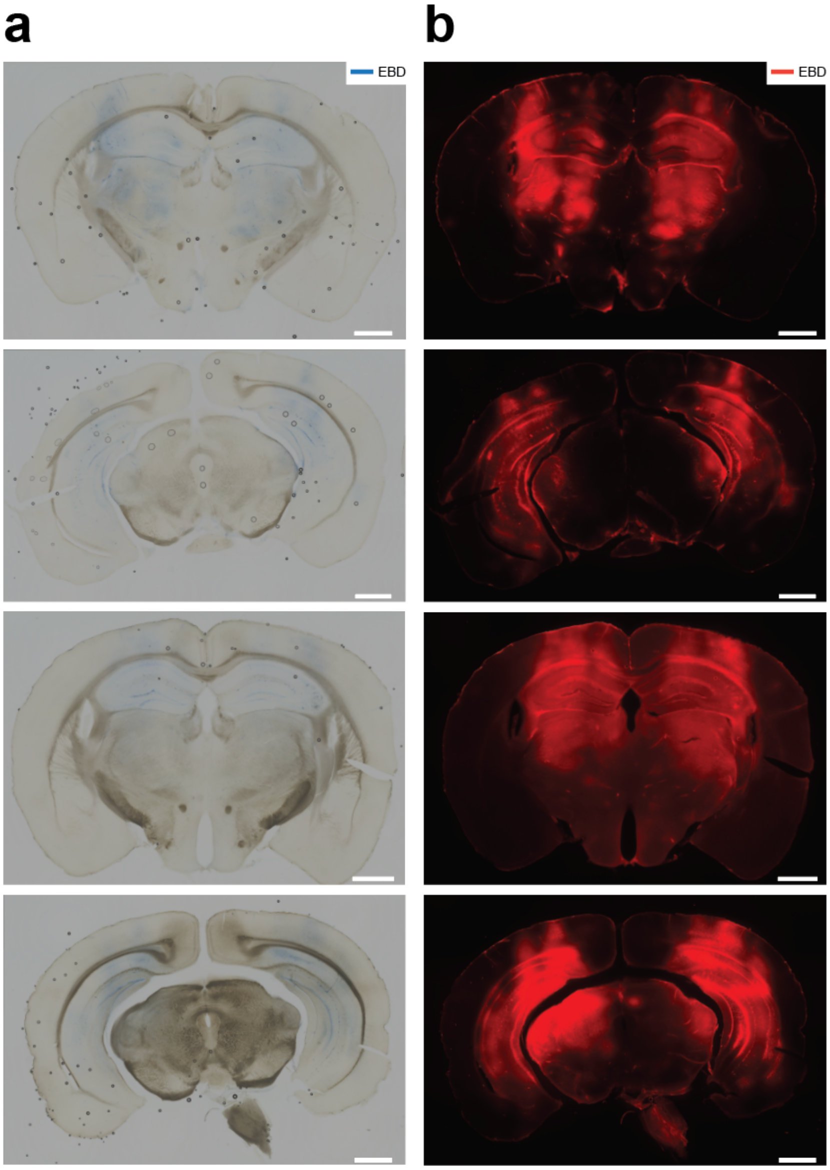


**Supplementary Fig. S2. Evaluation of FUS-BBBO targeting accuracy and map validation.** A total of four animals were insonated with FUS and injected with EBD. Each animal was insonated bilaterally with 12 FUS beams on the hippocampus of each hemisphere. a, The bright field image of the represented mouse for both dorsal (upper panel) and ventral (bottom panel) hippocampi. b, EBD fluorescence of the same brain section. Scale bar, 1mm. No obvious massive apparent tissue damage was seen in the sections.

**Supplementary Fig. S3.**

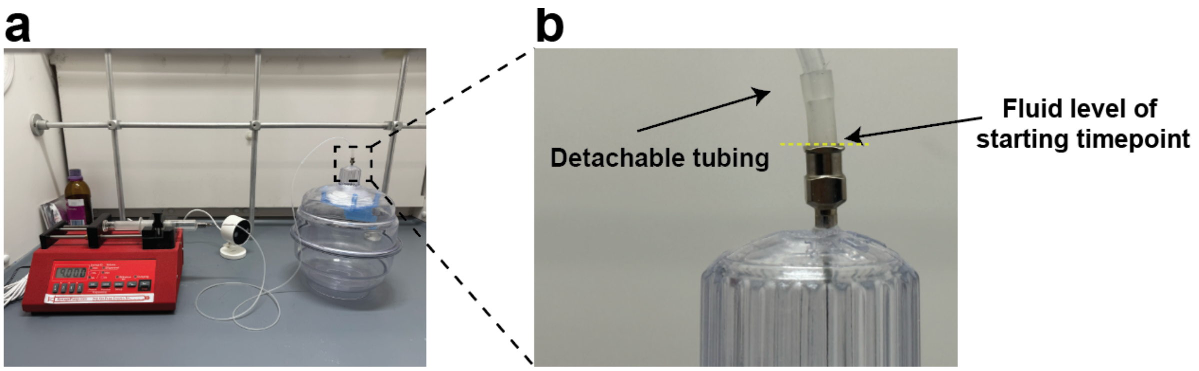


**Supplementary Fig. S3. Overview of Flurothyl Seizure Induction Setup.** a. Image of the flurothyl infusion and induction system. The setup consists of a syringe filled with 10% flurothyl connected to tubing, which is attached to an 18-G needle positioned at the top of the induction chamber. b. A closer view of the connection interface with detachable tubing. The dashed line indicates the flurothyl level as the starting point of the latency measurement.

**Supplementary Fig. S4**

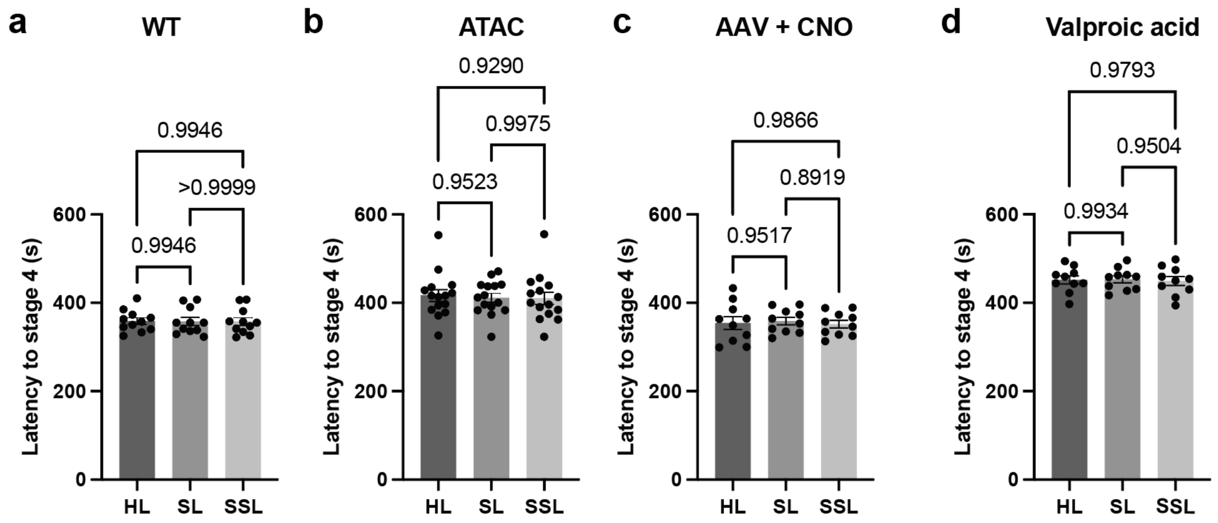


**Supplementary Fig. S4. Interexperimenter variability for flurothyl-induced generalized seizure latency across different experimental groups.** Video was reviewed and timed by two other blinded researchers to determine the latency to stage four seizure severity on a modified Racine scale. Groups are compared to each other using a Tukey honestly significant difference post hoc test. a, Latency of wildtype animals (F (2, 30) = 0.006549). b, Latency of ATAC-treated animals (F (2, 42) = 0.07579). c, Latency of AAV + CNO infused with CNO injected animals (F (2, 27) = 0.1077). d, Latency of valproic acid treated animals (F (2, 27) = 0.04750). We found no differences across all the groups among the three viewers (P > 0.8919 for all groups).

**Supplementary Fig. S5**

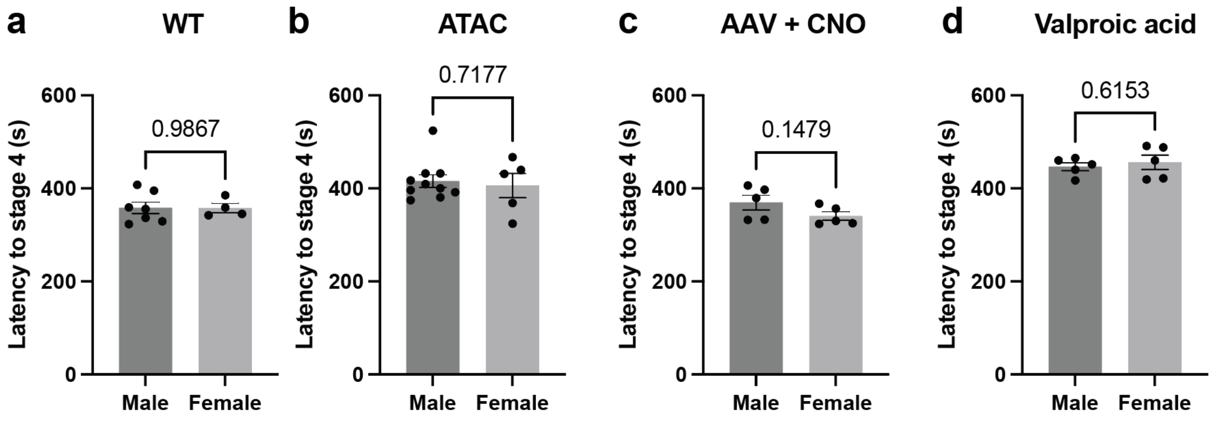


**Supplementary Fig. S5. Flurothyl seizure susceptibility in different sexes.** Sex differences were examined by separating the male and female animals from each group after a video review. A two-tailed t-test with unequal variance was used to compare two datasets. a, Susceptibility difference in wildtype animals (P = 0.9867). b, In ATAC-treated animals (P = 0.7177). c, In AAV + CNO injected animals (P = 0.1479). d. VPA treated animals (P = 0.6153). We found no susceptibility to a seizure caused by sex differences across all the groups in our flurothyl seizure model (P > 0.05).

**Supplementary table**

Supplementary Table 1. Results of the one-way ANOVA with Dunnett HSD post-hoc test comparing susceptibility of transfection of different fields of hippocampus with FUS-BBBO compared to a negative control, untargeted thalamus. v – ventral and d – dorsal. N = 15 mice were tested for each condition.

| Hippocampus field | Mean Diff. | Adjusted P Value | Summary |
|---|---|---|---|
| dCTX | 2.675 | 0.9953 | ns |
| dCA1 | -23.70 | <0.0001 | **** |
| dCA2 | -48.05 | <0.0001 | **** |
| dCA3 | -44.06 | <0.0001 | **** |
| dDG | -16.22 | 0.0024 | ** |
| vCTX | 1.044 | >0.9999 | ns |
| vCA1 | -18.92 | 0.0002 | *** |
| vCA3 | -30.11 | <0.0001 | **** |
| vDG | -27.49 | <0.0001 | **** |
